# The Organelle in the Ointment: cryptic mitochondria account for many unknown sequences in cross-species microbiome comparisons

**DOI:** 10.1101/2021.02.23.431501

**Authors:** Dylan Sonett, Tanya Brown, Johan Bengtsson-Palme, Jacqueline L. Padilla-Gamiño, Jesse R. Zaneveld

**Affiliations:** School of Pharmacy, University of Washington, Seattle, Washington, USA; University of Washington, Bothell, School of Science, Technology, Engineering, and Mathematics, Division of Biological Sciences, Bothell, WA, USA; Division of Systems and Synthetic Biology, Department of Life Sciences, Chalmers University of Technology, Gothenburg, Sweden; Department of Infectious Diseases, Institute of Biomedicine, Sahlgrenska Academy, University of Gothenburg, Gothenburg, Sweden; Centre for Antibiotic Resistance Research (CARe) at the University of Gothenburg, Gothenburg, Sweden; University of Washington, School of Aquatic and Fisheries Sciences, Seattle, WA, USA

**Keywords:** microbiome analysis, animal microbiomes, mitochondrial diversity, amplicon sequencing, mitochondria

## Abstract

The genomes of mitochondria and chloroplasts contain ribosomal RNA (rRNA) genes, reflecting their evolutionary ancestry as free-living bacteria prior to endosymbiosis. In microbiome studies of animals, plants, or other eukaryotic hosts, these organellar rRNAs are often amplified. If identified, they can be discarded, merely reducing sequencing depth. However, incorrectly annotated mitochondrial reads may compromise statistical analysis by distorting relative abundances of free-living microbes. We quantified this by reanalyzing 7,459 samples from seven 16S rRNA sequencing studies, including the microbiomes of 927 unique animal genera. We find that under-annotation of cryptic mitochondrial reads affects multiple of these large-scale cross-species microbiome comparisons, and can be severe in some samples. It also varies between host species, potentially biasing cross-species microbiome comparisons. We propose a straightforward solution: by supplementing existing taxonomies with diverse mitochondrial rRNA sequences, we resolve up to 97% of unique unclassified sequences in some entire studies as mitochondrial (14% averaged across all studies), without increasing false positive annotations in mitochondria-free mock communities. Overall, improved annotation decreases the proportion of unknown sequences by ≥10-fold in 2,262 of 7,459 samples (30%), including representatives from 5 of 7 studies examined. While standard DADA2 analyses are severely affected, the default positive filter in Deblur run through QIIME2 discards many divergent mitochondrial sequences, preventing bias in analysis, but also making analysis of these sequences more difficult. We recommend leveraging mitochondrial sequence diversity to better identify, remove and analyze mitochondrial rRNA gene sequences in microbiome studies.

## Introduction

Endosymbiotic theory has amassed considerable evidence that the ancestors of all animal mitochondria were free-living alpha-proteobacteria, while chloroplasts derive from formerly free-living cyanobacteria. Traces of the evolutionary history of mitochondria and chloroplasts as formerly free-living microbes can be found in organelle genomes. For example, mitochondria encode their own version of the small subunit rRNA gene called the 12S rRNA. Such organellar rRNA genes can pose problems for microbiome analysis, because they are often amplified by the same PCR primers used in 16S rRNA studies of the microbiome. For example, in a study of the microbiome of 32 plant species, contamination by plastid rRNA genes accounted for 23% of reads on average, but in certain host taxonomic groups, that rose as high as 94%^1^. Some animal taxa, such as scleractinian corals, have been diversifying for far longer than angiosperm plants, suggesting that they too may show substantial species-to-species variation in organelle rRNA sequences. If so, it is essential to ensure that these diverse mitochondrial rRNA sequences are efficiently removed prior to microbiome analysis.

The issue of organelle sequences confounding 16S rRNA gene studies has been addressed by excluding organelle sequences using molecular or *in silico* methods. Several molecular methods for exclusion of organelle SSU rRNA gene sequences have been developed, including peptide-nucleic-acid (PNA) clamps^1^ and Crispr-Cas9 cleavage^2^. However, such methods must generally be adapted to each host separately based on the host’s mitochondrial rDNA sequence, which may make their application challenging in cross-species surveys such as the Sponge Microbiome Project ^3^ and Global Coral Microbiome Project^4^. Additionally, applying such methods adds time and complexity to analyses and cannot be applied retroactively to existing studies without reamplification and resequencing of the underlying samples. Application of different molecular mitochondrial removal protocols tailored to specific taxonomic groups may also have difficult to quantify effects on the comparability of diverse studies in meta-analysis.

An alternative approach is to identify and filter out organelle rRNA sequences *in silico* using standard taxonomy annotation pipelines such as the naive-Bayesian RDP classifier^5^ and alignment-based algorithms such as USEARCH^6^ and VSEARCH^7,8^. If this process is accurate and unbiased across categories of samples, then removal of mitochondrial 12S rRNA sequences reduces effective sequencing depth but does not otherwise compromise microbiome analysis. Application of such methods to animal microbiomes typically does identify some mitochondrial 12S rRNA gene sequences. However, the existing literature does not establish whether existing workflows annotate all mitochondrial 12S rRNA sequences, or if additional mitochondrial sequences might be present in samples but under-annotated.

The problem of mitochondrial sequence removal is made much more challenging if multiple types of mitochondrial sequences are present in a study, but each varies in abundance between samples (**Fig. 1a**). For example, mitochondrial reads from diverse sources in an animal’s diet may be present in animal guts, while diverse microbial eukaryotes (each with unique mitochondria) are commonly found on tropical corals. Because microbiome data are compositional^9,10^, failure to remove mitochondrial sequences can distort the apparent relative abundance of bacteria and archaea present in the samples (**Fig. 1b**). Worse, biased annotation and removal of mitochondrial reads can further distort relative abundances if mitochondrial rRNA sequence diversity causes some mitochondrial sequences to be annotated while others are not (**Fig. 1c**). A key goal, therefore, is to uniformly identify all mitochondrially-derived rRNA sequences, thus preventing these reads from biasing downstream analyses (**Fig. 1d**).

**Figure 1.**
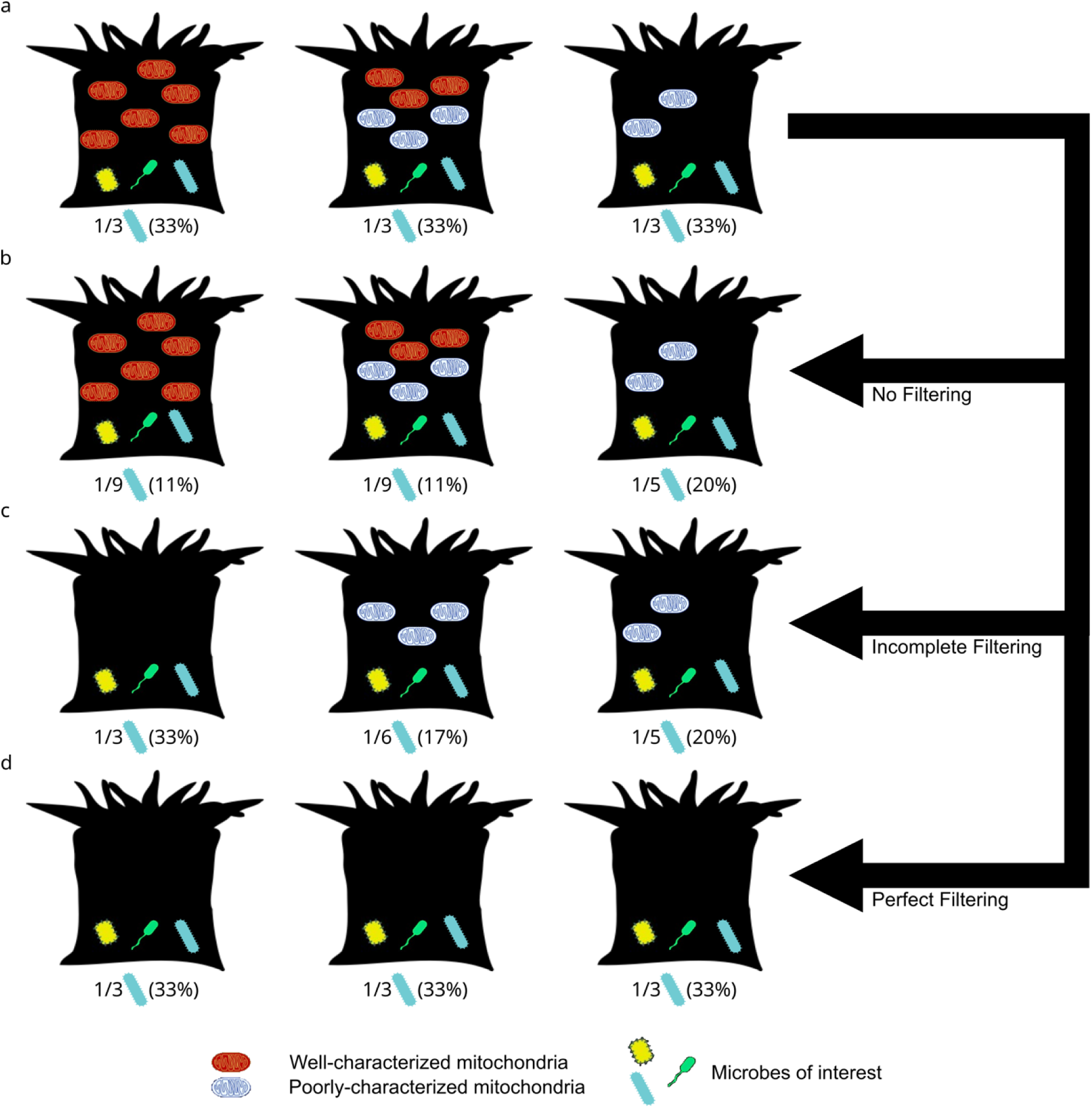
Conceptual diagram illustrating how taxonomically-biased misannotation of mitochondria can distort apparent relative abundances of microbes between animal species. **a** A set of 3 coral species with identical microbiomes, each of which has a microbe (blue microbe) with a true abundance of 33% (1/3 microbes) but with variation in abundance and type of mitochondrial rRNA sequences (red and blue mitochondria). **b** Analyzing microbiomes without removing mitochondria inflates the relative abundance of the microbe in samples with few mitochondria (3rd column). **c** Incomplete removal of mitochondrial sequences (red but not blue) distorts relative abundances based on both number and kind of mitochondria. **d** Perfect filtering of mitochondrial sequences removes mitochondrial abundance as a source of bias in diversity analysis.

In this manuscript we report widespread, severe, and host taxonomy specific underannotation of mitochondrial 12S rRNA sequences in several animal microbiomes when using standard taxonomy resources. Popular workflows typically identify some of the mitochondrially-derived reads in each sample. Surprisingly, however, some of these workflows do not necessarily annotate all, or even most mitochondrial sequences (similar to **Fig. 1c**). We demonstrate that this issue is taxonomically widespread. It severely affects analyses of the microbiome of reef-building corals, and to a lesser extent those of marine sponges, ants, birds and mammals. We develop an extended set of taxonomic annotations that are supplemented with diverse known mitochondrial 12S rRNA gene sequences, and demonstrate that this extended taxonomy resolves the provenance of the vast majority of ‘Unassigned’ sequences in some studies without causing false positive mitochondrial annotations.

## Results

### The mystery of high proportions of unknown bacteria in coral microbiomes

Coral microbiomes commonly report surprisingly high proportions of reads that cannot be assigned to a particular taxon, even at the domain level^11^. We encountered this issue in analyzing data on coral microbiome diversity as part of the Global Coral Microbiome Project (GCMP). This analysis collected DNA samples from phylogenetically diverse corals around the world (**Supplementary Table S1a**), and sequenced 16S rRNA gene amplicon libraries from them as part of the broader Earth Microbiome Project. In preliminary taxonomic analysis using QIIME 2^12^, we found that many samples showed extremely high proportions of microbes annotated as ‘Unassigned’ at the domain level (data not shown but replicated in later analysis; see below and **Supplementary Data Table 2a**).

In principle, these reads of ‘Unassigned’ taxonomy might represent novel diversity or sequencing artifacts. If these truly represented novel domain- or phylum-level diversity, that would be very surprising, given that such novel diversity has not appeared in studies of full-length 16S rRNA sequences from corals^13^, despite the identification of coral-specific members of several known phyla. Additionally, since the standardized sequencing methods used in the GCMP were also used in many other studies in the broader Earth Microbiome Project, it would be surprising if such high proportional abundances of ‘Unassigned’ reads were due purely to sequencing artifacts.

A third explanation is that unassigned sequences could represent under-annotated organelle rRNA sequences. We found that many of these ‘Unassigned’ reads had strong BLAST hits to known mitochondrial sequences of corals, algae, diatoms and other marine organisms (**Supplementary Data Table 3**), as well as potential contaminants (e.g. human). Of the 1,000 most abundant sequences in the GCMP dataset annotated as ‘Unassigned’ by VSEARCH using SILVA 138 as the taxonomic reference, 49.5% (1,342/2,713) of top 5 BLAST hits were mitochondrial (i.e. had ‘mitochondria’ in the description). Yet although some reads that showed high sequence similarity to mitochondria by BLAST were annotated as ‘mitochondria’ by VSEARCH, most were annotated as ‘Unassigned’ at the domain level. This persisted regardless of whether the SILVA or Greengenes database was used, despite substantial sequence similarity to known mitochondrial sequences. Therefore, we chose to explore the generality of this phenomenon, its effects on microbiome analysis, and potential solutions.

### Supplementation of taxonomic references with more diverse organelle rRNA sequences resolves many unknown sequences as mitochondria

If many ‘Unassigned’ rRNA reads do, in fact, represent mitochondria rather than sequencing artifacts or new domain-level diversity, we should expect that adding reference sequences for known mitochondrial rRNAs from diverse hosts should reduce ‘Unassigned’ annotations and increase mitochondrial annotations. Conversely, there is no reason to expect either sequence artifacts or microbes from novel lineages to show any special degree of sequence similarity with animal mitochondria. Thus, if either sequencing error or taxonomic novelty explained ‘Unassigned’ sequences, we should expect little change in sequence annotations when improving the diversity of mitochondrial sequences represented in reference taxonomies.

We quantified the number of mitochondrial and chloroplast reference sequences found in SILVA version 138^14,15^ and Greengenes 13_8^16^ (**Fig. 2, Supplementary Data Table 4**). We then collected additional mitochondrial and chloroplast rRNA gene sequences from Metaxa2^17^, and generated extended taxonomic references by integrating them into either SILVA 138 or Greengenes 13_8. This greatly expanded the number of mitochondrial and chloroplast sequences in each reference (**Fig. 2**). The additional sequences increased the number of mitochondrial sequences in SILVA from 420 to 3,799 (approximately 9-fold; **Fig. 2**), and the number in Greengenes 13_8 from 211 to 3,600 (approx. 16-fold). Chloroplast sequence supplementation increased Chloroplast rRNA diversity by 2.8-fold in SILVA 138, or 5-fold for Greengenes 13_8 (**Fig. 2**).

**Figure 2.**
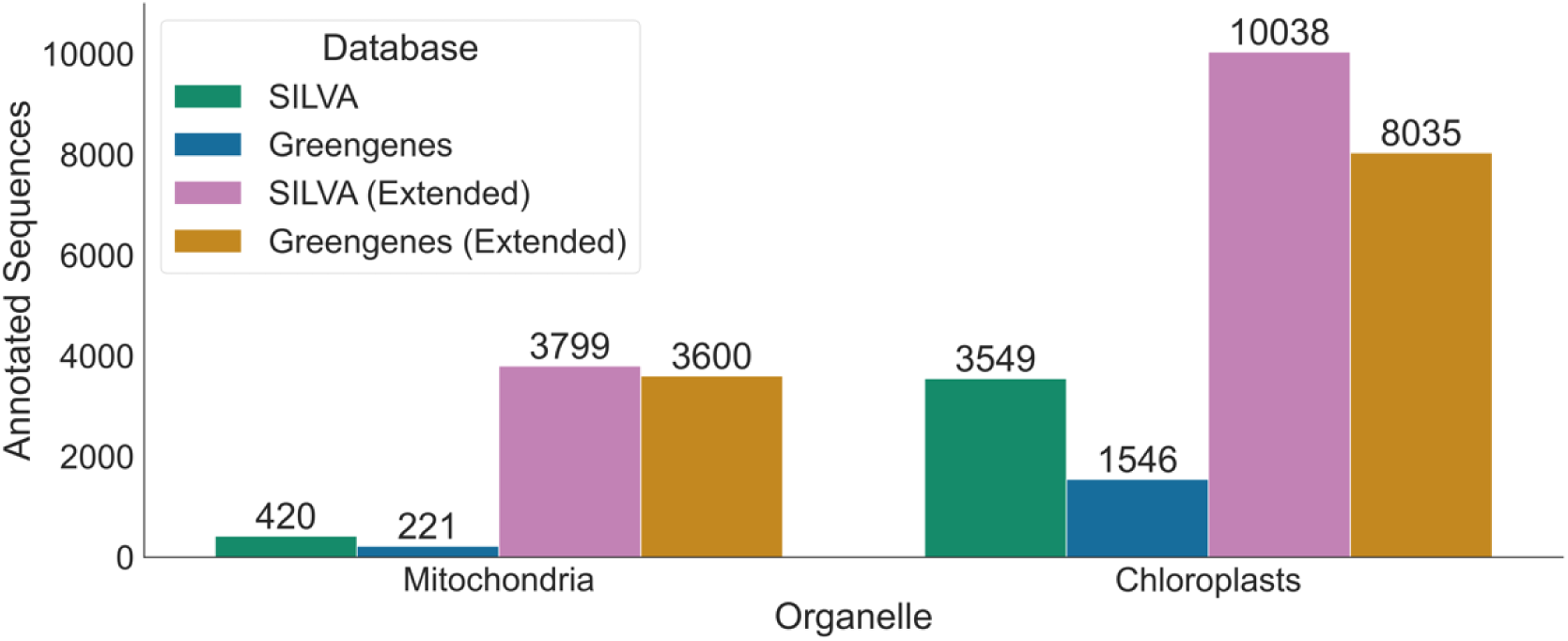
Counts of sequences annotated as mitochondria or chloroplasts in each reference taxonomy. SILVA refers to the SILVA 138 release; Greengenes to the Greengenes 13_8 release. Extended reference databases incorporate the original database in addition to organelle sequences from the Metaxa2 database.

We tested how adding these additional mitochondrial reference sequences affected mitochondrial annotation when using different combinations of denoisers (Deblur^18^ or DADA2^19^), base taxonomic references (SILVA^14,15^ or Greengenes^16^), and taxonomic classification methods (VSEARCH^7^ or naive Bayes^20^). We applied these tests to multiple datasets (**Supplementary Data Table 2a-e)**. These included data from the human microbiome^21^ and milk microbiomes^22^, as well as multiple cross-species surveys^3,4,23,24^ within animal groups (including ants^24^, marine corals^4^, marine sponges^3^, and other diverse vertebrates^23^).

### Effects of extending reference taxonomies differ across studies and animal groups

Addition of diverse reference mitochondrial sequences had very large effects on analysis of diverse animal groups (**Fig. 3**) for which proportionally few sequenced genomes are available (e.g. marine corals and sponges), but little effect in several single-species studies of well characterized animal hosts (e.g. in human microbiomes). Importantly, when expanding reference taxonomies decreased ‘Unassigned’ annotations (**Fig. 3a**), it typically also increased mitochondrial annotations (**Fig. 3b)**, consistent with many ‘Unassigned’ reads representing cryptic organelle sequences, rather than sequencing artifacts or novel diversity. Adding additional chloroplast diversity to taxonomic references also modestly increased chloroplast annotations (**Fig. 3c**) in studies that included herbivores (e.g. diverse birds and mammals), although these changes were minor compared to shifts in mitochondrial sequence annotation.

**Figure 3.**
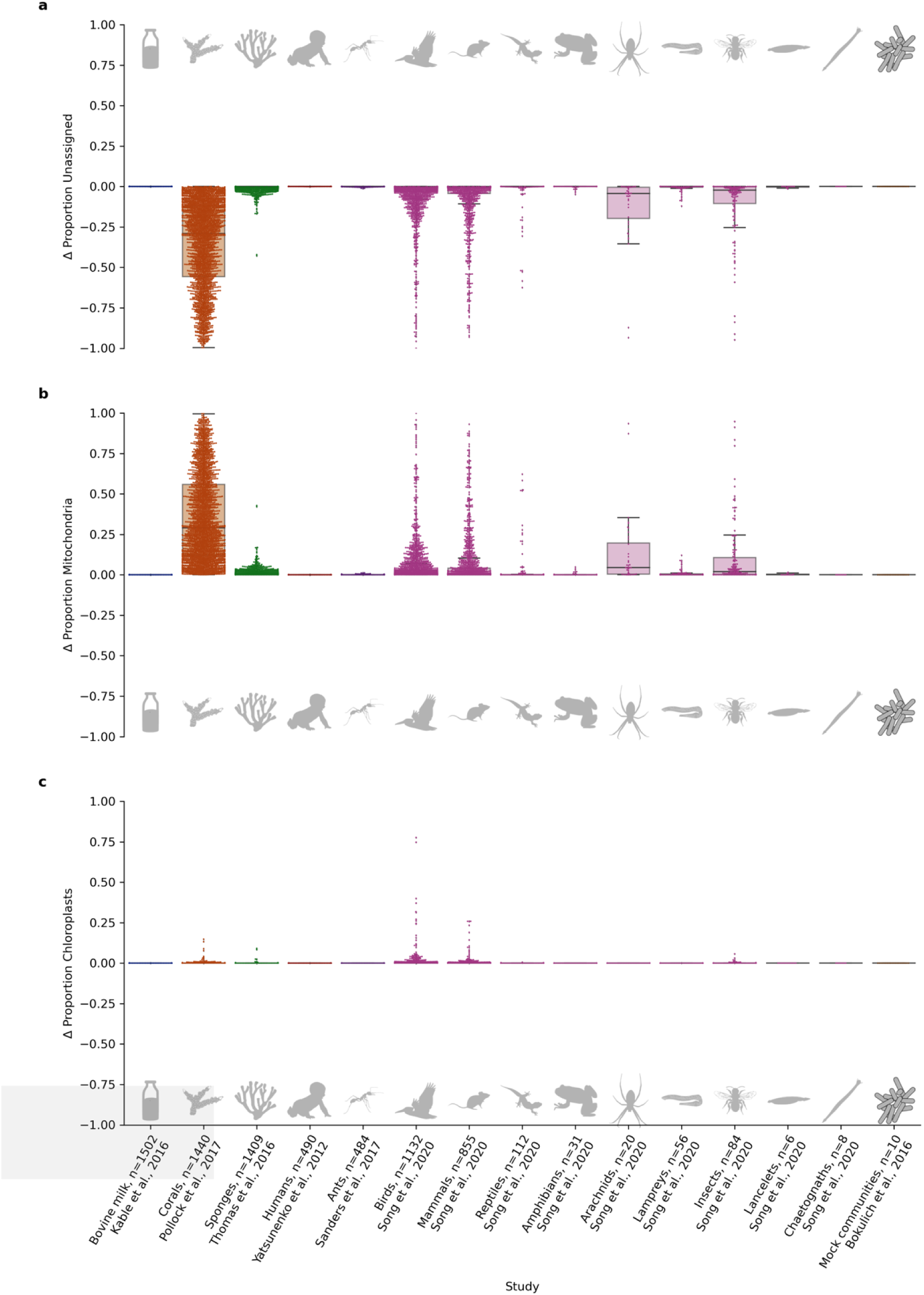
Supplementation of SILVA resolves many “Unassigned” microbes as mitochondria. Differences in apparent relative abundance when 16S rRNA gene sequencing data from several studies were re-annotated using a version of SILVA 138 with additional mitochondrial references (Methods). Annotation with the extended reference taxonomy decreases the proportion of unknown sequences by 10-fold or greater in 2262 of 7459 samples (30%), including representatives from 5 of 7 studies examined (71%), and these decreases were largely matched with proportionate increases in mitochondrial annotations. **a.** Difference in the proportion of reads that were unassigned (e.g. “Unassigned” at domain level). **b.** Difference in the proportion of reads that were classified as mitochondria. **c.** Difference in the proportion of reads that were classified as chloroplasts. Study labels include the clade studied, author, and number of samples.

Examining differential annotations confirmed that the vast majority of reannotations at the class level or above were formerly ‘Unassigned’ sequences reassigned as mitochondrial (93%) or chloroplast (6.5%) sequences (**Fig. 4**, **Supplementary Data Table 5c**). The only other trend notable in these reassignments was that at finer levels of taxonomic resolution some annotations shifted in their specificity (e.g. from species to genus level identification of some Firmicutes; **Supplementary Data Table 5g**). Notably, independent benchmarks of taxonomic analysis from 16S rRNA data using mock communities showing overconfident results below the family level^20^, suggesting that small shifts towards more conservative annotations in some cases are unlikely to obscure useful biological patterns.

**Figure 4.**
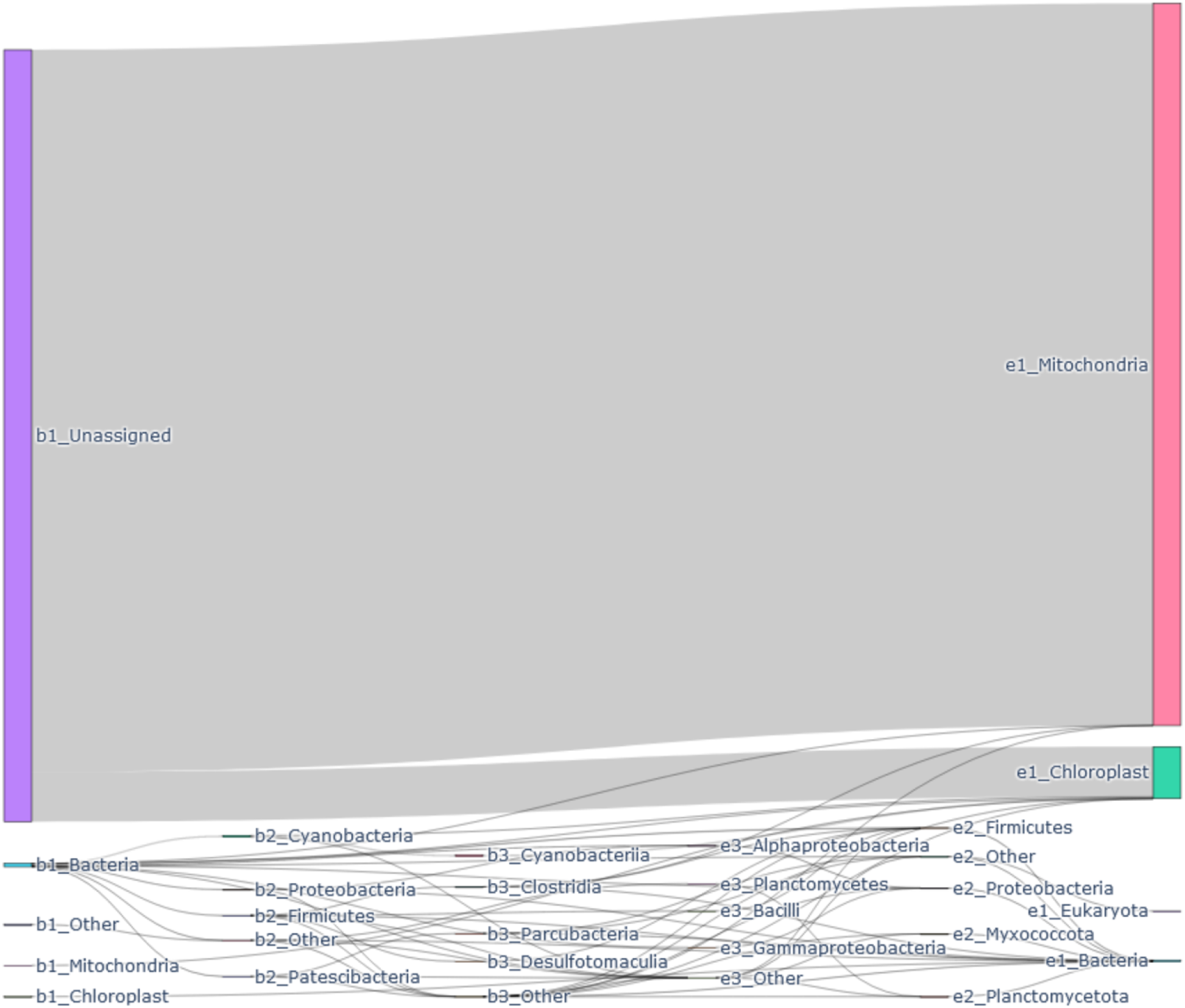
Reclassification of sequences using the extended SILVA taxonomy. Sankey diagram flows show differences in sequence classification between base (left) and extended (right) SILVA 138 taxonomy at or above the class level, weighted by total sequence frequency across all studies. Wider bars indicate reclassification of either many ASVs or a smaller number of ASVs with high frequencies. Prefixes indicate classification under the base (b) or extended (e) taxonomies and taxonomic rank (1, Domain; 2, Phylum; 3, Class). For clarity of visualization (this figure only), mitochondria and chloroplasts are displayed at the top level of the taxonomy, rather than within Alphaproteobacteria or Cyanobacteria, and taxon nodes not in the top four most frequent of each level were collapsed to “Other”. The most common alterations were of Unassigned sequences reassigned to Mitochondria (∼5.2 million out of 5.6 million total altered annotations, 93.1%), followed by Unassigned reads being reassigned to Chloroplast (∼360,000 / 5.6 million annotations, 6.5%). No Unassigned sequences were reannotated as non-organelle Bacteria, Archaea, or Eukaryota. All other reassignments account for less than 0.5%, with the most common non-organelle reassignment reannotating unassigned Bacteria to Firmicutes (3618 out of 5.6 million, 0.07%).

### A positive filter against known 16S rRNA sequences also prevents mitochondrial contamination

The default Deblur pipeline implemented in QIIME 2 includes a ‘positive filtering’ step. In this step, sequences are filtered against the Greengenes 88% OTU reference taxonomy. Those that do not fall within a 65% sequence identity threshold and 50% coverage threshold to this reference database are removed. Thus this positive filtering step demands that sequences broadly resemble known 16S rRNA sequences of free-living bacteria or archaea, or reference organelle sequences present in Greengenes. The threshold was selected to incorporate the range of known variation in bacterial and archaeal 16S rRNA sequences across phyla. If under-annotated mitochondrial reads are divergent, then this step may explain the better performance of deblur vs. DADA2 with default settings. To test this, we denoised sequences while either adding a positive filter to DADA2 or suppressing the default positive filter used in deblur (**Fig. 5**) and then annotated the results. By manipulating the positive filtering step in this way, we traced differences in mitochondrial annotation between deblur and DADA2 to the positive filtering step used in the QIIME2 implementation of deblur.

**Figure 5.**
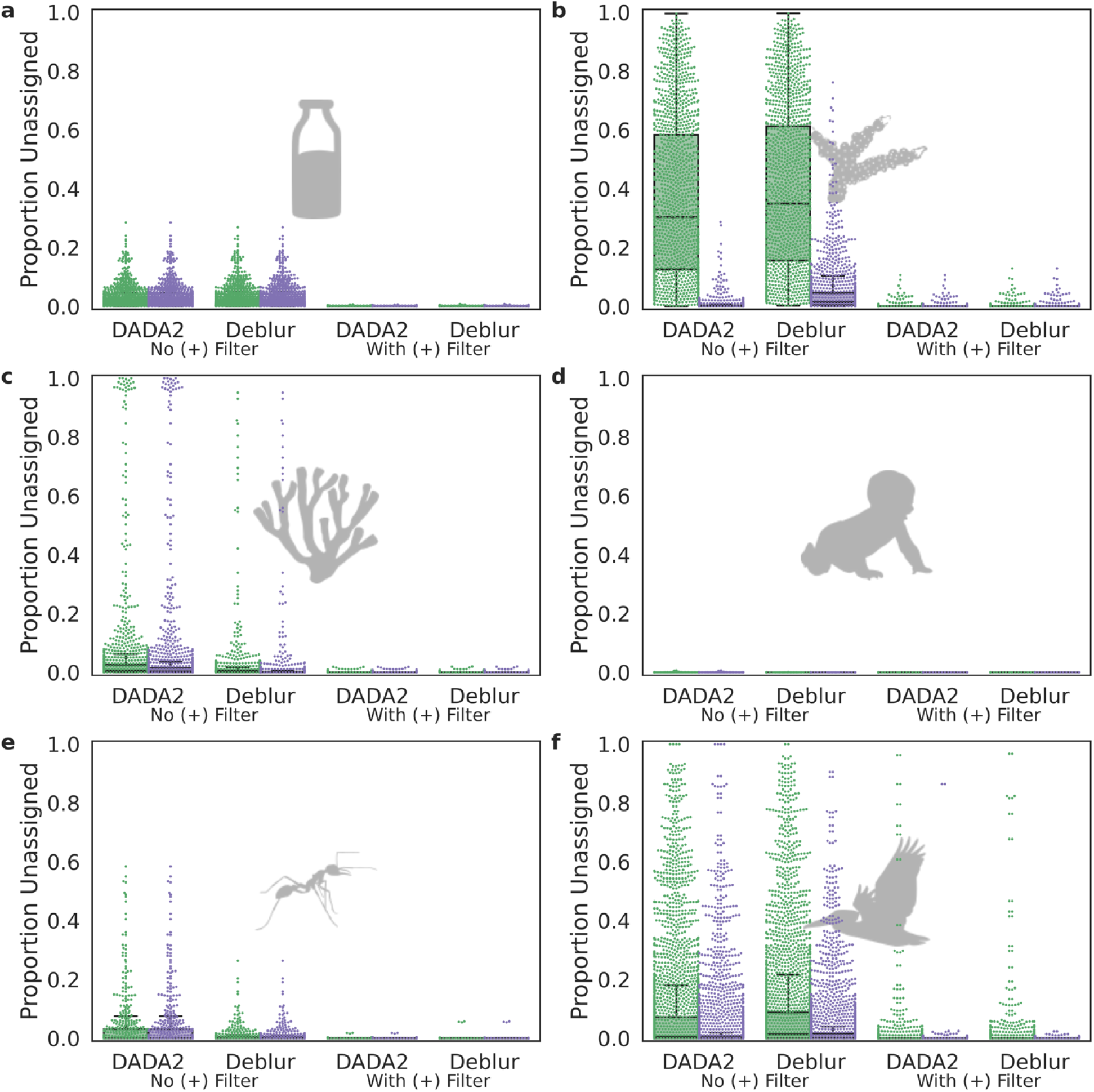
The presence of a positive filter explains much of the difference between Deblur and DADA2. To determine the cause of the substantial difference in “Unassigned” annotations when using different denoising methods, we separately investigated the methods and the SortMeRNA positive filter generally applied to the QIIME2 implementation of Deblur. Using the filter (gray shading) severely reduced differences in the proportion of unassigned sequences across denoising methods and base vs. extended SILVA reference taxonomies, relative to the unfiltered (unshaded) samples (excepting **d.** human gut samples from Yatsunenko *et al*. in which samples were extremely well-characterized). **a.** Bovine milk samples from Kable *et al*. **b.** Coral samples from Pollock *et al*. **c.** Marine sponge samples from Thomas *et al*. **d.** Human gut samples from Yatsunenko *et al*. **e.** Ant gut samples from Sanders *et al*. **f.** Diverse vertebrate samples (all classes) from Song *et al*.

The cryptic mitochondrial or chloroplast reads detected when using an expanded rather than base taxonomy seem to overlap heavily with divergent sequences excluded by Deblur’s positive filter. Adding an identical SortMeRNA^25^ positive filtering step as is used in Deblur to DADA2 effectively eliminates the differences in how DADA2 and Deblur respond to the expanded taxonomies. This is likely because adding the SortMeRNA positive filter to the DADA2 workflow causes cryptic mitochondrial reads to be filtered out, meaning that the expanded taxonomy no longer changes the results much. Conversely, suppressing the positive filter from the Deblur workflow causes the expanded taxonomies to matter much more than they otherwise would (**Fig. 5**). However, even with a positive filter, the expanded taxonomies seem to influence mitochondrial annotations in some samples. For example, expanded taxonomies reduced the number of samples with high levels of Unclassified sequences in the Song *et al.* dataset of diverse vertebrate microbiomes, even when a positive filter was present (**Fig. 5**).

### Expanded mitochondrial reference taxonomies do not promote false positive annotations

A potential concern about expanding reference taxonomies with extra mitochondrial sequences (some of which are lower in quality than average for Greengenes or SILVA) is that it might lead to false positive annotations of mitochondrial taxonomy. We used two approaches to test for this. First, we annotated the taxonomy of microbial communities of known composition (mock communities^8,19,26–29)^ using either standard or expanded taxonomies. Since these mock communities were constructed without real mitochondria, we treated any mitochondrial annotations as false positives. However, the expanded taxonomies did not increase mitochondrial annotations in these mock communities (**Fig. 3b**).

Another hypothesis we considered was that the expanded taxonomies might cause increased false positive annotations of sequencing artifacts as mitochondria (perhaps due to increased incidental matches in nucleotide sequences). We tested this by scrambling the sequences from the GCMP coral dataset as well as sequences from the mock communities known to lack mitochondria. We expected the annotation for such scrambled sequences to be “Unassigned”, since they should retain no non-random sequence similarity to known rRNA genes (sharing only their mononucleotide frequencies). After generating scrambled rRNA sequences, we then re-annotated these sequences using either base or expanded taxonomies, and attributed any increase in mitochondrial annotations as evidence of either signal from mononucleotide frequencies (in the GCMP dataset only) or false positives (possible in either the mock or GCMP datasets equally). When using the VSEARCH classifier, which relies on sequence alignment, no sequences were annotated as mitochondria or chloroplasts, but rather were all Unassigned at the domain level (**Fig. 6a, 6c**).

**Figure 6.**
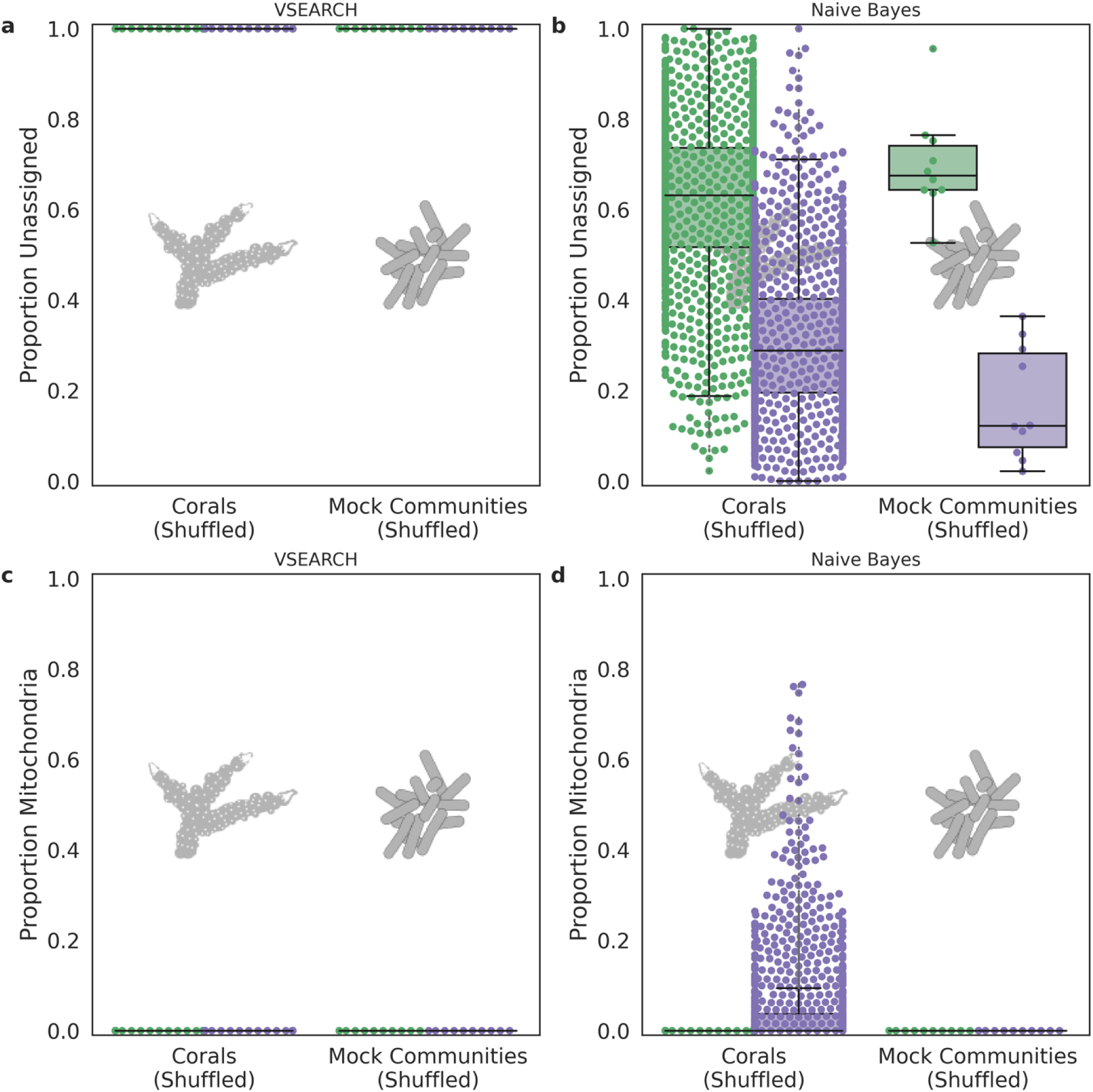
Expanded reference databases do not increase false positive annotations of simulated sequencing artifacts. To test whether sequencing artifacts not removed by denoising might be classified as mitochondria when using expanded reference taxonomies, we scrambled sequences of the GCMP (expected to contain mitochondria) and mock datasets (expected to contain no mitochondria), and used the VSEARCH (**a**, **c**) or naive Bayes (**b**, **d**) classifier to assign taxonomic annotations according the the base SILVA reference taxonomy (green). This procedure was repeated for the expanded taxonomy (purple), with the distribution of “Mitochondria” annotations plotted on the y-axis. Using the different reference taxonomies, the VSEARCH classifier showed no change in annotations of mitochondria. However, the naive Bayesian classifier annotated substantially fewer sequences as ‘Unassigned’ in each shuffled dataset, driven by an increase in Bacteria annotations (with no assignment at the phylum level) in both datasets, and an increase in Mitochondria annotations in the GCMP.

When using the naive Bayes classifier, more scrambled sequences from the GCMP dataset were annotated as mitochondrial (4.5% vs. 0.001%) when using the extended rather than the base version of SILVA, with a corresponding change in ‘Unassigned’ annotations (**Fig. 6b, 6d**). This was somewhat surprising — scrambling the sequences destroys all information other than nucleotide frequencies — and we expected any mitochondrial annotations in scrambled sequences to reflect increased false positives. However, we also tested the possibility that this machine learning method is using raw mononucleotide frequencies (which are not changed by scrambling) to identify mitochondrial reads.

If differences in mitochondrial annotation when using expanded taxonomies were driven by false positives, we should expect them to also appear when the naive Bayes classifier is applied to mock community data known not to contain mitochondria. However, in that case, we saw no difference in the number of mitochondria annotated with base vs. extended taxonomic references using either VSEARCH or the naive Bayes classifier (**Fig. 6c, 6d**). This suggests that diversifying mitochondrial sequences in reference taxonomies does not increase false positive mitochondrial annotations. Interestingly, the expanded taxonomies do change annotations of some previously unassigned non-mitochondrial sequences, reannotating sequences unassigned at the domain level to unclassified bacteria (**Fig. 6c**).

### Under-annotation of mitochondrial reads can influence alpha and beta-diversity

We sought to understand whether changes in taxonomic annotation caused by mitochondrial under-annotation could influence comparisons of microbiome alpha or beta diversity. To do so, we reran select alpha and beta diversity analyses for each study after using different mitochondrial removal methods and compared the results (**Fig. 7**). For example, we compared 1) the influence of age for human samples, 2) season on milk microbiomes, and 3) the family-level taxonomy of the host for ant, marine sponge, and coral microbiomes. These specific categories were chosen based on the reported results of each study. The results indicate that most differences in comparisons of alpha and beta diversity are more subtle than for taxonomic analysis. We compared differences across categorical variables in each study using four alpha diversity metrics (Faith’s phylogenetic diversity, **Fig. 7a**; observed features, **Fig. 7b**; Shannon diversity, **Fig. 7c**; Simpson’s evenness measure E, **Fig. 7d**) and four beta diversity measures (Unweighted UniFrac distance, **Fig. 7e**; Weighted UniFrac distance, **Fig. 7f**; Jaccard distance, **Fig. 7g**; Bray-Curtis dissimilarity, **Fig. 7h**). Reassuringly, overall differences in effect size were modest, ranging from 0.85-fold to 1.09-fold. Generally these changes in effect size were greater for qualitative, presence-absence based beta diversity measures (e.g. Jaccard distance) than for quantitative ones (Weighted UniFrac or Bray-Curtis).

**Figure 7.**
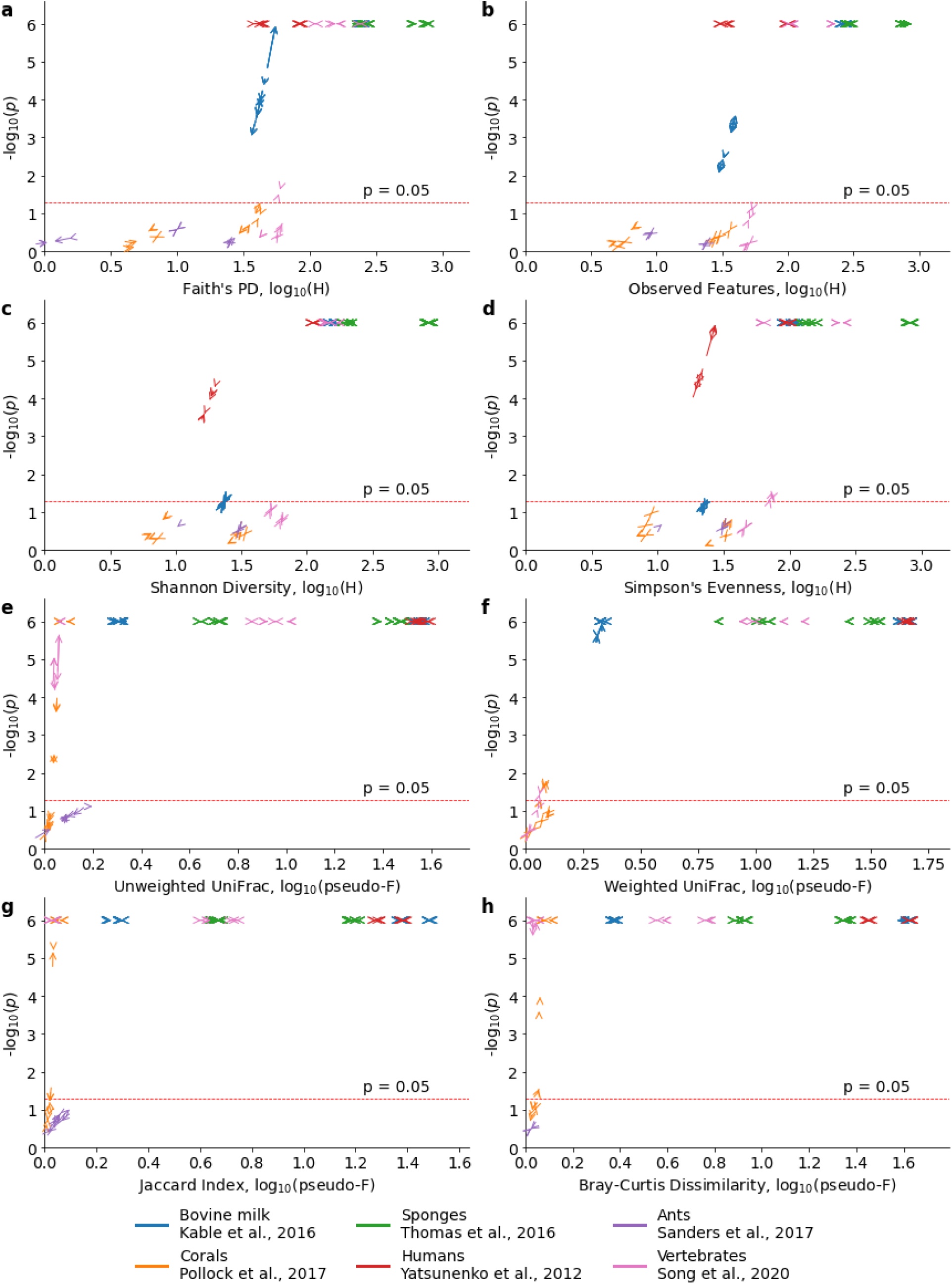
Change in effect size and significance between base and extended taxonomy references. Changes in taxonomic classification and filtering can affect both the effect size and significance of Kruskal-Wallis and PERMANOVA tests comparing alpha (**a**, Faith’s Phylogenetic Diversity; **b**, count of observed features; **c**, Shannon Diversity Index; **d**, Simpson’s Evenness) and beta diversity (**e**, Unweighted UniFrac; **f**, Weighted UniFrac; **g**, Jaccard Index; **h**, Bray-Curtis Dissimilarity) across metadata categories. Arrow plots show the effect of using the extended taxonomy on study-specific tests (see **Supplementary Data Table S7** for a list of metadata categories tested and statistical results). Alpha diversity p-values are capped at 10^-6^ for clarity of visualization, while beta diversity p-values are limited by the PERMANOVA permutations (10^6^). The dashed line represents p = 0.05. Multiple comparisons may necessitate adjusting this cutoff.

Nonetheless, in several cases these modest changes in effect size were sufficient to alter p-values, including one case which would change the significance of the test (**Fig. 7; Supplementary Data Tables 7a-d**). Across all comparisons, the mean absolute shift in p-values by annotation was 0.006, with more comparisons shifted downward in p-value when the extended taxonomy was used. One interpretation is that improved identification and removal of mitochondrially-derived reads can somewhat improve the ability to detect biologically interesting trends in host-associated microbiomes.

While most biological questions would be answered the same way regardless of mitochondrial removal protocol, these changes are sufficient to change biological conclusions in some cases. We identify 25 combinations of study, category, denoiser, filter, and alpha or beta diversity metric in which mitochondrial annotation method differences would be sufficient to shift nominal p-values from below the (arbitrary) p = 0.05 threshold to above it, or above p=0.05 to below it (**Supplementary Data Tables 7b, 7c**). Of comparisons where mitochondrial annotation method mattered, the extended taxonomies resulted in 7 nominal p values that became “not significant” (at alpha = 0.05), and 18 p-values that became “significant”. These examples include whether coral mucus, tissue and skeleton have distinct microbial communities as assessed by the Jaccard distance; whether coral families differ in richness as assessed by Faith’s Phylogenetic diversity; whether milk storage silos significantly differ in Shannon diversity or Simpson’s evenness of their milk microbiome communities; and whether vertebrate classes (i.e. Mammalia vs. Reptilia) differ in gut microbiome evenness (Simpson’s evenness).

We recognize that in practice many studies must correct for multiple comparisons, so the true range of p-values that impact biological conclusions will vary by study design. While the range of possible methods and degrees of multiple comparison vary greatly, we identify 53 cases in which the change in p-values between mitochondrial annotation methods was greater than ±0.05. Finally, as many studies emphasize effect sizes and their 95% confidence intervals rather than significance *per se*, we also examined how mitochondrial annotation method impacted effect sizes. On average, the annotation method did not substantially change effect sizes (mean fold-change in effect size 0.998). However, we identify 5 cases where effect sizes changed by 2-fold or more, all of which were comparisons involving whether Peruvian ants differed in gut microbiome richness or evenness by habitat (**Supplementary Data Table 6d**).

We note that in order to standardize this analysis we ran each study through a common pipeline, so a difference in our analysis does not necessarily mean that the study conclusions themselves are suspect. For example, in several cases Deblur pipelines were used that resolve these issues, but our benchmarks suggest that issues could have been encountered if DADA2 without additional filtering steps or an extended mitochondrial reference taxonomy had been used.

## Discussion

Microbiome studies have become vital tools in medicine, ecology and evolution. However, best practices for many aspects of marker gene studies of microbiomes continue to develop. In this study we focus on the effects of different methods for annotation of organelle rRNA sequences, and their potential to influence biological conclusions.

### Cryptic mitochondria bias microbiome analysis

In comparing samples that contain different mitochondrial sequences (including many cross-species comparisons), we find that differences in the accuracy with which mitochondrial reads are identified by taxonomic annotation pipelines can impact apparent microbial relative abundances, as well as community properties like alpha and beta diversity.

In cases where only a single mitochondrial sequence is present in each sample, it may be easy to detect if mitochondrial annotation has failed, because no reads will be annotated as mitochondrial. Investigators could then take *ad hoc* steps to remove mitochondrially-derived sequences. However, there are several mechanisms by which multiple types of mitochondrially-derived sequences may be present in 16S rRNA gene samples. For example, if the tissues of dietary, parasitic, or epiphytic organisms are co-mingled with the focal organism in samples, it can result in diverse mitochondria that must be annotated. Additionally, some animals and many plants show considerable heteroplasmy^30^ in which mitochondrial genome sequence varies within the same individual. Levels of intra-individual sequence divergence between mitochondria can be substantial (e.g. up to 23% divergence reported in lobster mitochondrial 12S rRNAs^31^). Transposition of mitochondrial DNA to the nucleus, which is common (e.g. in humans^32^) can generate nuclear mitochondrial sequences (NUMTs). These are known to confound eDNA studies, and may also be amplified in 16S rRNA gene studies.

If any of these mechanisms are in operation, it is possible to annotate some but not all mitochondrially-derived sequences, offering researchers a false sense of confidence that all sequences have been correctly identified. In light of the results presented in this manuscript, it appears common for one or more of these mechanisms to create situations where only some mitochondrially-derived sequences in a 16S rRNA are correctly annotated. The approaches described here provide additional security against distortions due to cryptic mitochondrially-derived sequences when any of these common situations occur.

### Present benefits and future opportunities for improved mitochondrial annotations

Cross-species microbiome comparisons, such as the GCMP, the Sponge Microbiome Project, and Song *et al.*, often must identify many host species in the field during sample collection. In cases where this is challenging, correctly annotated mitochondrial sequences may offer clues. For example, the mitochondrial reads in the 16S data of the GCMP conflicted with the initial field identifications of several coral samples by divers. The identification of the coral species was subsequently revised based on the combined evidence provided by this molecular data and reexamination of sample photographs.

Laboratories may also be able to use analysis of mitochondrial sequences to detect and identify sources of contamination. For example, both human and bird mitochondrial sequences were detected in a small number of GCMP samples when unknown sequences were queried with BLAST (**Supplementary Data Table 3**), suggesting some samples that might be contaminated with non-host DNA during sampling or sequencing, and could be excluded from analysis.

### Recommendations

Our results suggest several actionable steps that can be taken for cross-species microbiome comparisons. First, researchers should be aware that high proportions of unknown sequences may be attributable (among other causes) to cryptic organelle rRNA sequences, both of the host organism, and any dietary or symbiotic eukaryotes. Second, by supplementing standard reference taxonomies with diverse mitochondrial sequences, as described here, researchers can in many cases greatly improve annotation of cryptic organelle sequences. Third, if such additions are not used, a positive filter against known rRNA sequences can remove divergent organelle sequences (as well as the sequencing artifacts that such positive filters were designed to exclude). Fourth, users of online repositories of marker gene data, including Qiita^33^, should take care to check whether either a positive filter or an expanded reference taxonomy has been applied. Fifth, cross-species comparisons of microbiome diversity should carefully implement all of these precautions, since the relative abundance of cryptic mitochondrial reads can vary across species. Finally, maintainers of taxonomic reference databases should take special efforts to include diverse mitochondrial and chloroplast sequences, as well as nuclear sequences derived from them (i.e. NUMTs homologous to mitochondrial rRNA genes) and recognize that diversity in these organelle sequences can be just as important as diversity within bacterial groups for correct annotation of amplicon data.

## Conclusion

Together, these results demonstrate that cryptic mitochondrial sequences are important confounders in cross-species microbiome comparisons. Successful annotation and subsequent removal of these sequences depends on the diversity of organelle sequences in reference taxonomies, and on whether a positive filter is used to exclude divergent sequences. These results further confirm that extending reference libraries to account for mitochondrial rRNA diversity expands mitochondrial annotations without causing an increase in false positive annotations, regardless of whether VSEARCH or naive Bayes classifiers are used. Therefore, we recommend expanding the diversity of organelle rRNA sequences in taxonomic references. Consistent mitochondrial annotations will both help prevent bias in microbiome analyses, and also provide important contextual information about studies, such as the presence of contamination from non-target samples.

## Materials and Methods

### Workflow code

Analyses were conducted using Jupyter notebooks, python scripts, and shell scripts. Unless noted otherwise, default parameters were used for each analysis step. The full set of code is publicly available on GitHub (https://github.com/zaneveld/organelle_removal).

### Initial generation of the Global Coral Microbiome Project dataset

Coral microbiome DNA sequences were selected from samples collected by the GCMP as described in Pollock *et al*., but including additional locations outside of Australia in Panama, Saudi Arabia, Columbia, Singapore, and Réunion that were not described in that manuscript. Importantly, these samples have been sequenced twice: once using Illumina (Illumina, Inc., San Diego, California, USA) MiSeq sequencing, and again using the EMP protocol and Illumina HiSeq sequencing. The samples analyzed here were sequenced following the EMP protocol, as this was the larger sample set, and also used standardized methods applied to diverse study systems.

Briefly, samples were collected from water, sediment, and the mucus, tissue, and skeleton of corals from 457 coral colonies, then DNA was extracted using the MoBio Powersoil DNA Isolation Kit (MoBio Laboratories, Carlsbad, California, USA) and processed by the Earth Microbiome Project at the Center for Microbiome Innovation (University of California San Diego, San Diego, California, USA). PCR was run on the V4 region of the 16S rRNA gene using 515f/806r primers (5’-GTGTGCCAGCMGCCGCGGTAA-3’) and sequenced using Illumina HiSeq with 125bp paired-end reads. Sequences were downloaded from the Earth Microbiome Project via Qiita project ID 10895 (specifically prep id 3439). In Qiita, these sequences were processed using standard EMP workflows: fastq files were demultiplexed using 12bp Golay codes with the QIIME 1.9.1 split_libraries script (default parameters), trimmed to 100nt, and then subjected to quality control with deblur 1.1.0 (default parameters). The “deblur final table” artifact (ID: 59201, now deprecated) was used for initial investigations.

### Initial detection of high numbers of mitochondria annotated as “Unassigned” in the GCMP dataset

We initially discovered many samples in the GCMP dataset with high proportions of reads which were labeled “Unassigned” by the QIIME2 feature-classifier plugin^20^ using the classify-consensus-vsearch method^7^ and the Greengenes 13_8 reference taxonomy.

We queried the 1,000 highest frequency “Unassigned’’ sequences with blastn against the nt database (blast_unknowns.py), with the following options: ‘-max_target_seqs 5’, ‘-max_hsps 1’, ‘-outfmt 6 qseqid sseqid staxids stitle evalue bitscore’. **(Supplementary Data Table 3).**

### Study selection

To determine if under-annotation of mitochondrial reads was widespread across multiple studies, we reanalyzed the original sequences from five additional studies in the Qiita database, covering sponges^3^, diverse vertebrates^23^, humans^21^, bovine milk^22^, and ants^24^. These studies were selected to represent a range of animal-associated study systems in which we expected mitochondrial sequences to be present. Samples from mockrobiota^8,19, 26–29^, a collection of artificially constructed (mock) microbial communities which were known to lack mitochondria, were used as negative controls.

### Construction of expanded SILVA and Greengenes databases

We downloaded and extracted reads from the Metaxa2^17^ custom BLAST database. Mitochondrial reads in this database were themselves curated from Mitozoa (version 2.0, release 10)^34^ and SILVA (release 111)^15^. Chloroplast sequences were collected from the Phytoref database^35^. Sequences annotated as mitochondria in the Metaxa2 reference database were inserted into the SILVA and Greengenes databases, along with the Phytoref sequences. In all cases, taxonomy strings were reformatted to match SILVA or Greengenes conventions, respectively. This resulted in new custom databases which we refer to as “extended” reference taxonomies (e.g. Greengenes (Extended)).

### Initial data processing and quality control

We downloaded the raw upstream fastq files of each study from Qiita and imported them into QIIME 2 2021-4. After demultiplexing (q2-demux emp-single^36,37^), sequences from Yatsuenko et al. were converted to Phred33 from Phred64 by exporting and reimporting into QIIME 2. Each study was separately denoised with DADA2 (q2-dada2) and Deblur (q2-deblur). To isolate the effect of the SortMeRNA positive filter against the Greengenes 13_8 88% OTUs included by default in QIIME 2’s implementation of Deblur, we ran each denoising step with and without the filter applied (**Supplementary Figure S1**).

### Annotation and Benchmarking Workflow

Sequences were classified using the QIIME2 feature-classifier^20^ plugin with the classify-consensus-vsearch^7^ and classify-sklearn^38^ methods, with base and extended reference taxonomies. Taxa counts were generated with q2-taxa barplot, after which mitochondria and chloroplasts were filtered from the feature tables (q2-feature-table filter-features). Samples in each table were rarefied to 1000 sequences, after which samples not present in every feature table of each study were discarded to allow for direct comparison of composition and diversity across studies and methods.

### Testing the effects of deblur’s SortMeRNA positive filter step

We noticed a substantial difference in annotations of ‘Unknown’ and ‘Mitochondria’ between denoising algorithms (DADA2 vs Deblur) when we explored the effect of supplemented reference taxonomies (**Supplementary Data Table S3**). In QIIME2, the Deblur plugin uses a ‘positive filter’ step in which SortMeRNA^25^ is used to search sequences against a reference 16S rRNA database (Greengenes 13_8 88% OTUs by default), discarding any sequences below 65% identity with 50% coverage to at least one reference sequence. This filtering step is not present when DADA2 is used. We hypothesized that the SortMeRNA positive filter — rather than algorithms themselves — might be responsible for the differences. We tested this using two analyses: first, we ran a version of deblur in which we disabled the SortMeRNA positive filter by inputting reference sequences matching each query, causing all sequences to avoid the filter. The results of this procedure are labeled ‘deblur_unfiltered’, in contrast to the default, positively filtered Deblur results, which we refer to as ‘deblur_filtered’. Second, we added a SortMeRNA filtering step after denoising with DADA2, using settings identical to the Deblur SortMeRNA defaults. This tested whether addition of a positive filtering step would be sufficient to bring the levels of mitochondrial rRNA exclusion for DADA2 in line with that of deblur. We call default DADA2 results ‘dada2_unfiltered’, and those with a SortMeRNA positive filter added ‘dada2_filtered’. Across these four categories of results (deblur_unfiltered’, deblur_filtered, dada2_unfiltered, and dada2_filtered), we compared the proportion of unassigned reads (**Fig. 5**)

### Testing changes to annotation of *in-silico* scrambled sequences

To further validate that extending the SILVA/Greengenes taxonomies did not increase false positive mitochondrial annotations, we generated shuffled versions of the GCMP data (high proportion of unknown sequences) and mock community data (zero unknown sequences). In these shuffled datasets, true biological sequences were scrambled at the mono-nucleotide level. We reasoned that — aside from nucleotide composition — this procedure would destroy any biological signal in the sequences. Thus, any differential annotation of these sequences would be attributable to either a) the minimal biological signal conveyed in nucleotide frequencies or b) false positive annotations.

### Testing changes in diversity analysis

Under-annotation of mitochondrial reads has the potential to alter alpha or beta diversity estimates, especially when under-annotation varies between sample categories (e.g. if different host species are being compared). To quantify these effects, we took samples from each study (considered separately) and compared alpha and beta diversity results when comparing the SILVA vs. SILVA (Extended) reference taxonomies. We tested all combinations of denoiser (Deblur vs. DADA2), classifier (VSEARCH vs. naive Bayes), and filtering (presence or absence of a positive SortMeRNA filter). Within these datasets, we selected two metadata categories per study for comparison of alpha and beta diversity. These were anatomy (’tissue_compartment’) and family-level taxonomy (’family’) for Pollock *et al*., species (‘host scientific name’) and sample type (‘empo3’) for Thomas *et al*. life stage (‘life_stage’) and environment (‘env_biome’) for Yatsunenko *et al*., ‘season’ and silo (‘silo_lot_id’) for Kable *et al*, ‘genus’ and ‘habitat’ for Sanders *et al*., and ‘class’ and ‘country’ for Song *et al*.

## Acknowledgements

The authors would like to acknowledge Daniel McDonald, Nicholas Bokulich, Justin Shaffer for useful discussions. This work was supported by a NSF IOS CAREER award (#1942647) to J.Z.

## Supplementary Information

### SI Figures

**Figure S1.**
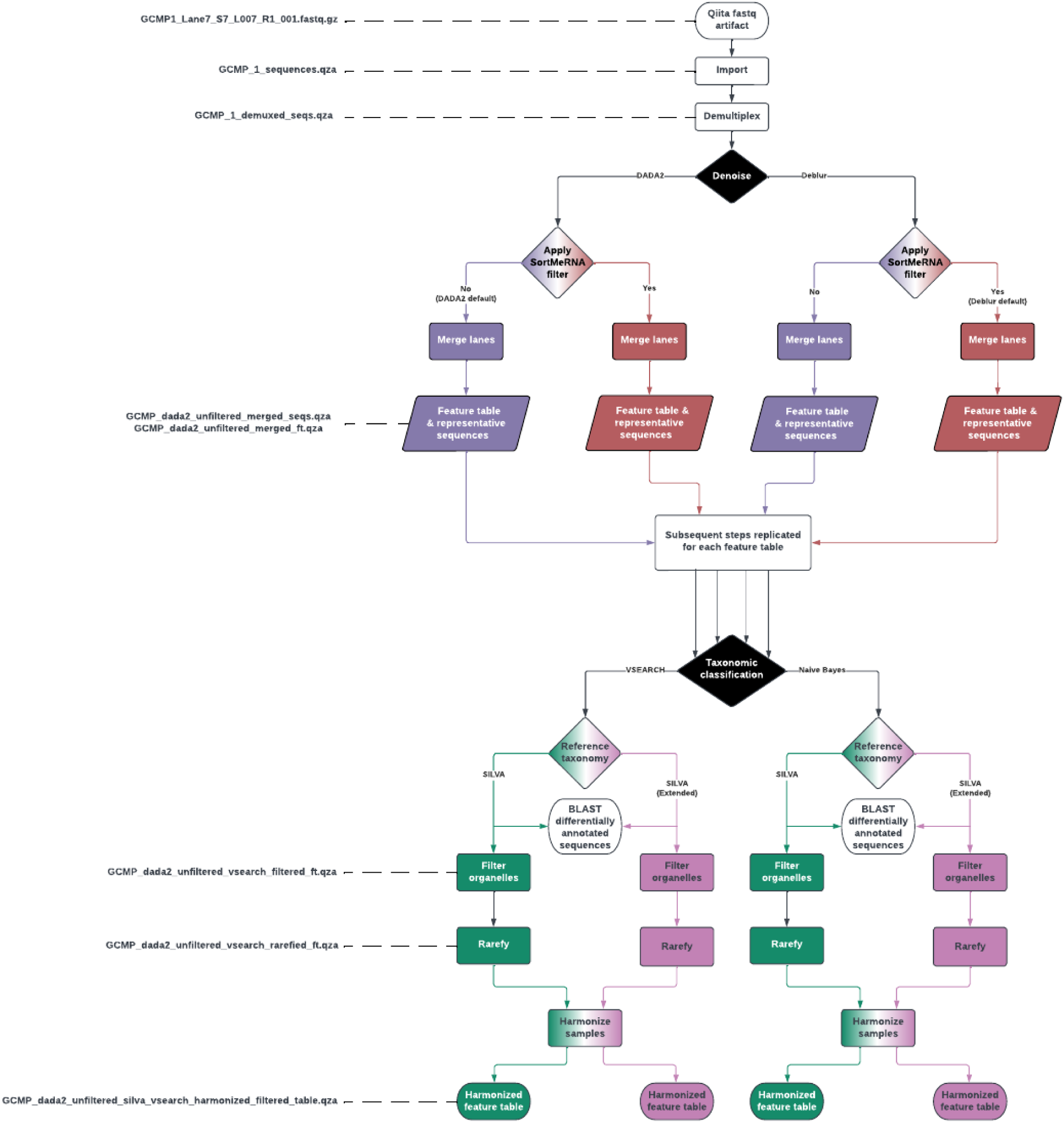
Workflow diagram for analyses in the study. Sequence files for each study in the analyses were downloaded from QIITA (6/7 studies) or mockrobiota (mock community samples). They were then denoised using either deblur or DADA2, with or without a positive filtering step against known sequences (this is default for deblur but not DADA2; see text). For each of the resulting 28 feature tables were run through taxonomic classification with either VSEARCH or a naive Bayes classifier from scikit-learn (as implemented in QIIME2), and with either SILVA or our extended SILVA reference taxonomy, resulting in 112 feature tables. Sequences identified as deriving from organelles were removed, each sample was rarefied to 1000 sequences per sample (discarding samples with fewer than 1000 reads). Finally, to enable fair comparison between rarefied base and rarefied extended taxonomies, only the intersection of samples from these tables was analyzed. Additionally, all the above steps were repeated on Greengenes 13_8 (Supplemental Results)

### SI Data Tables

**Table S1.** Metadata for all studies in the analysis. **a.** Metadata for Pollock *et al*., the Global Coral Microbiome Project **b.** Metadata for Thomas *et al*., the Sponge Microbiome Project **c.** Metadata for Yatsunenko *et al*., study of human gut microbiomes. **d.** Metadata for Kable *et al*., study of bovine milk microbiomes **e.** Metadata for Sanders *et al*., study of Peruvian ant microbiomes. **f.** Metadata for Song *et al*., meta-analysis of vertebrate microbiomes

**Table S2.** Effects of adding mitochondrial sequence diversity to taxonomic references. **a.** Proportion and absolute numbers of reads annotated as Unknown, Mitochondria or Chloroplast. Reports a summary of annotations for all combinations of denoisers, base or extended reference taxonomies, and classification methods by sample. **b.** Five number statistical summary of the proportion or absolute number of reads annotated as Unknown, mitochondria or chloroplast by source study. This is a summary of the data in S2a, giving the minimum, first quartile, median, third quartile and maximum. **c.** Mann-Whitney U tests of proportions of samples annotated as ‘Unassigned’, ‘Mitochondria’, or ‘Chloroplast’ comparing base reference taxonomies to their extended counterparts.

**Table S3.** Results of BLAST searches for top 1000 most abundant Unknown sequences identified in the Global Coral Microbiome Project dataset

**Table S4.** Counts of organelle sequences in Greengenes 13_8 and SILVA 138 before and after supplementation with additional sequences.

**Table S5.** Counts of sequences across all studies which were annotated differently by base SILVA 138 and SILVA (Extended) using VSEARCH. **a.** Domain-level reannotations. **b.** Phylum-level reannotations. **c.** Class-level reannotations. **d.** Order-level reannotations. **e.** Family-level reannotations. **f.** Genus-level reannotations. **g.** Species-level reannotations.

**Table S6.** Results of negative control analysis testing for annotation of mitochondria in scrambled GCMP sequences using base or reference taxonomies.

**Table S7.** Impact of mitochondrial annotation reference on alpha and beta diversity test statistics for **a.** all tests examined, **b.** tests that shifted from p < 0.05 to p > 0.05 when using extended taxonomic references. **c.** tests that shifted from p > 0.05 to p < 0.05 when using extended taxonomic references. **d.** tests with a > 2-fold difference in effect size when using extended taxonomic references.

### Supplementary Data Files

**Supplementary Data File 1.** Extended version of the SILVA 138 FASTA file, containing original SILVA sequences and additional mitochondria and chloroplast sequences. Along with Supplementary Data File 2, this allows for annotating the taxonomy of 16S rRNA gene amplicon sequences (using QIIME2 or other microbiome software packages) according to the ‘extended’ taxonomies described in this manuscript.

**Supplementary Data File 2.** Extended version of the SILVA 138 taxonomy file, mapping both original and extended sequences from the FASTA file to taxonomic annotations. Along with Supplementary Data File 1, this allows for annotating the taxonomy of 16S rRNA gene amplicon sequences (using QIIME2 or other microbiome software packages) according to the ‘extended’ taxonomies described in this manuscript.

